# Immobilised enzyme reactors for post-production glycan modification of purified glycoproteins

**DOI:** 10.64898/2026.07.09.737398

**Authors:** Nicholas J. DeBono, Joel Cain, Chi-Hung Lin, Nicolle H. Packer, Edward S.X. Moh

**Author notes:** **Correspondence:** Dr Edward S.X. Moh **, Contact address:**.

## Abstract

Controlling protein glycosylation as a critical quality attribute of biopharmaceuticals remains challenging when glycosylation is coupled to cellular production systems. Here, we present a proof-of-concept glycosyltransferase immobilised enzyme reactor (IMER) housed within a 3D-printed column that enables directed post-production glycan modification of purified glycoproteins. Using β-1,4-galactosyltransferase (β4GalT1-IMER) and α-2,6-sialyltransferase (ST6Gal1-IMER) immobilised on Ni-NTA resin, the IMER achieved near-complete galactosylation and substantial sialylation of partially deglycosylated bovine fetuin N-glycans with their respective substrates with a maximum substrate-enzyme contact time of four minutes. Isomeric-level analysis revealed arm-specific addition preferences for both enzymes, consistent with known specificities. The modular IMER design permits sequential connection of individual enzyme chambers, potentially offering a scalable, plug-and-play platform for constructing defined glycan structures on recombinant glycoprotein therapeutics.

## Introduction

Protein glycosylation is a well-established critical quality attribute (CQA) of biopharmaceutical products (1-3). This is due to the critical role glycans have in structural variability and receptor interactions (4), fundamentally changing how a glycoprotein interacts with its surrounding environment. Due to this classification as a CQA, glycosylation is targeted as an attribute for control to ensure ongoing safety and efficacy of existing glycoprotein therapeutics, particularly given the dominance of monoclonal antibodies in the current biologics market (5). The ability to control glycosylation is therefore of great interest for ensuring ongoing pharmaceutical CQA control, particularly if glycoengineering methods can increase the yield of active therapeutic glyco-products. Glycoengineering methods have been extensively reviewed previously(6-9), both in an *in vivo* and *in vitro* setting.

The industrial production of glycoprotein therapeutics relies predominantly on mammalian cell lines, particularly Chinese Hamster Ovary (CHO) cells (10, 11). While these systems give high titres, the glycans they install are not always appropriate for the target therapeutic, and non-native features can interfere with essential interactions (for example, Neu5Gc eliciting immunogenic responses (12)). Glycosylation of a given protein varies greatly between expression hosts (13), and as a CQA this variation can exclude otherwise attractive production systems, including ‘generally recognised as safe’ (GRAS) bacterial and fungal hosts (14). Non-mammalian hosts such as insect, plant and yeast cells (7) instead produce simpler or partially complete glycans. Rather than a limitation, this partial glycosylation provides a defined starting substrate that can be matched to a downstream finishing step, in which specific monosaccharides are added to the glycans already present on a purified glycoprotein.

If protein glycosylation could be decoupled from other existing cellular processes, the concern of off-target glycosylation producing unwanted interactions can be alleviated. This can be achieved through creation of a rudimentary synthetic Golgi apparatus, used after protein production and purification from any recombinant production host. Suitable production hosts for producing ‘partially complete’ humanised glycosylation for finalisation in an external Golgi apparatus are currently commercially available(15), and existing glycans can also be trimmed through exoglycosidase treatment. Decoupling glycosylation enables the use of production systems that are less than ideal in terms of glycosylation but may be a preferable option when compared to other metrics such as titre of yield, growth rate, strain robustness and cost of strain maintenance(16). A theoretical synthetic Golgi apparatus has been proposed (17) and partially realised (18), with a specific focus on producing monoclonal antibodies. The implementation of a post-production, artificial Golgi can assist with the maintenance of the CQAs of glycosylation within the defined parameters required for a wider range of recombinantly produced glycoproteins.

An artificial Golgi can be realised with an immobilised enzyme reactor (IMER), where enzymes are fixed to a solid support either through physical containment or chemical conjugation(19). The enzymes in an IMER are reusable and have the potential for much higher throughput in a continuous flow system when compared to free enzymes, as well as having compatibility with downstream analytical techniques such as HPLC. Kinetically, immobilised enzymes may be subject to a less than ideal environment as they are bound to an immobile phase(20), however as glycosyltransferases are type II membrane proteins and anchored in the Golgi, immobilisation through N-terminal domain can more closely mimic their natural environment, making them an ideal target for an IMER. The application of glycosyltransferases in a continuous-flow system also reduces potential potent by-product inhibition by physically removing products as they are made. The SUGAR-TARGET platform immobilises glycosyltransferases on beads to galactosylate IgGs but runs as discrete batch reactions, separating the glycoprotein from the enzymes between steps (18), while the microgel-based MiRAGE reactor operates under continuous flow but assembles free oligosaccharides rather than modifying glycans already present on a purified glycoprotein (21). As yet, no system combines continuous flow with direct modification of the existing glycans on an intact, purified glycoprotein.

Here we develop and characterise two glycosyltransferase IMERS, β-1,4-galactosyltransferase (B4GalT1) and α-2,6-sialyltransferase (ST6Gal1) immobilised via 8xHis tag to Ni-NTA affinity within a 3D-printed housing in a continuous flow setting with a glycoprotein substrate. Using de-sialylated and/or de-galactosylated bovine fetuin as a model N-glycoprotein substrate, we demonstrate near-complete galactosylation on the B4GalT1-IMER and substantial α2,6-sialylation ST6Gal1-IMER within ∼160 s of substrate-enzyme contact time. Monitoring the same column performance at time points over multiple hours gives an idea of IMER (and enzyme) reusability. The modular, plug-and-play chamber design offers a scalable platform for post-production glycan finishing of recombinant glycoprotein therapeutics.

## Materials and Methods

### 3D modelling and printing of the IMER

All models of the IMER were developed using Fusion 360 (version 2.0.17721 x86_64) (Supplementary Figure 1). Models were printed on a Formlabs Form 2 SLA printer with Formlabs Clear Resin v4 at 0.4 µm layer height and 100 % infill. After printing, models were sonicated in acetonitrile for 15 minutes, then cured for 1 hour in a custom 495 nm UV lightbox. After curing, the exterior of each model was coated with clear nail polish to improve the translucency of the plastic.

### Glycosyltransferase production by HEK293 cells

Mammalian expression vectors containing genes coding for human β-1,4-galactosyltransferase (pGEn2-B4GalT1) and human α-2,6-sialyltransferase (pGEn2-ST6Gal1) on pGEn2 vector were purchased from the DNASU plasmid repository (22). Both vectors contain a coding region for an N-terminus 8xHis/Strep purification tag followed by a Green Fluorescent Protein (GFP) fusion linker and soluble domain of the enzymes.

HEK293 were maintained in FreeStyle 293 Expression Medium (Thermo Fisher) in a shaking incubator (Eppendorf) at 37°C and supplied with 5% CO_2_. For enzyme expression, HEK293 were seeded at 2.5 x10^6^ cells/mL in 50 mL FreeStyle 293 Expression Medium in a 125 mL baffled cell culture flask (Corning). 150 µg of pGEn2-ST6Gal1 or pGEn2-B4GalT1 were prepared at 0.5 µg/µL in FreeStyle 293 Expression Medium and added dropwise into the cell culture. The flask was incubated in the shaking incubator for 5 minutes. 450 µg of PEI was prepared at 0.5 µg/µL in FreeStyle 293 Expression Medium and added dropwise into the culture. The culture was put back to the shaking incubator and cultured for 24 hours. 0.5 mL of

220 mM valproic acid (VPA) stock solution was added into culture to final concentration of 2.2 mM VPA. The culture was incubated for another 48 hours before enzyme purification. Culture was centrifuged at 500 g for 5 minutes and supernatant was collected for purification.

### Glycosyltransferase purification and loading of the IMER

Freshly charged nickel-nitrilotriacetic acid (Ni-NTA) 0.4 µm agarose resin beads were used for enzyme immobilisation and purification (ThermoFisher, cat # 88221). These resin beads were equilibrated in HEPES-HCl pH 7.4 containing 150 mM NaCl after stripping nickel ions with EDTA and charging with nickel acetate. 1 mL of equilibrated resin was added to culture media from transfected HEK293 cells containing secreted 8xHis-Strep tagged glycosyltransferases, spiked with 10 mM imidazole and 150 mM of NaCl to reduce non-specific binding. The supernatant containing the resin beads was rocked for an hour at 4 °C, then protein bound resin was collected by sedimentation and pipetted into a 3D printed column previously plugged with wetted cotton wool at one end to prevent resin beads from flowing through. Columns were then capped and stored at 4 °C until required for use.

### Preparation of substrate and reaction components for IMER

Bovine fetuin was treated with either α2-3,6,8 Neuraminidase (NEB) to remove terminal sialic acids, or with α2-3,6,8 Neuraminidase and β-1,4-galactosidase (NEB) to remove terminal sialic acids and then terminal galactose saccharides, by incubating the fetuin with the enzymes in the presence of GlycoBuffer II (NEB) at 37 °C for 3 hours according to manufacturer’s recommended enzymatic concentrations. Effective de-sialylation and de-galactosylation of the fetuin glycoprotein substrate, and activity of recombinantly expressed ST6Gal1 and B4GalT1 was confirmed with SDS-PAGE (Supplementary S2, S3).

### Galactosylation and sialylation using the IMER

All IMER column experiments were performed with a running buffer of HEPES-HCl pH 7.4, 10 mM MnCl_2_ and 150 mM NaCl. Reaction mixtures for each experiment thus contained final concentrations of 10 mM MnCl_2_, 150 mM NaCl, 100 nmols of appropriate donor sugars (either UDP-galactose or CMP-N-acetylneuraminic acid depending on glycosyltransferase used) and appropriately pre-treated partially deglycosylated fetuin (20 µg). Packed IMER components were connected and placed in the column oven of an Agilent 1260 HPLC and equilibrated to 37 °C before injection of the reaction mixture. Injections were made at 0, 2, 4 and 8 hours, with each injection run at both 25 µL/min and 75 µL/min so that the two flow rates could be compared at every timepoint. Between injections, the column was kept within the column oven at a flow rate of 5 µL/min. After each injection, fractions were collected directly into a PVDF membrane 96-well plate using the fraction collector module. Samples were then taken for N-glycan analysis by MS.

### Released N-glycan analysis of IMER products

N-glycans were directly released in duplicate from the protein product fractions on the PVDF membrane as previously described (24, 25) and analysed by LC-MS/MS. Briefly, PVDF protein spots from the IMER eluent were blocked with 1% (v/v) polyvinylpyrrolidone in water, washed twice with water, and left to dry. Dried discs were then reactivated with ethanol, and 5 units of PNGase F (NEB) in 15 µL ultra-pure water was added to each sample, which was subsequently incubated overnight at 37 °C.

Released N-glycans were collected and then deamidated with 10 mM ammonium acetate pH 5, for 1 hr before drying under vacuum. N-glycans were then reduced using 1 M NaBH_4_ in 50 mM KOH at 50 °C for 3 hr, diluted with 100 µL ultrapure water, then neutralized with 2 μL glacial acetic acid. Self-packed 10 μL tips containing carbon (Supelclean ENVI-Carb) (approximately 15 mm packing height) on a C18 stage tip were first washed with 50 μL of 90% acetonitrile (ACN) and 0.1% trifluoroacetic acid (TFA) and equilibrated twice using 50 μL of 0.1% TFA. The diluted glycan samples were applied onto the column in two loadings of 60 μL, washed with water, and eluted using 50 μL of 50% ACN and 0.1% TFA. Eluted glycans were then dried using a vacuum concentrator.

### PGC-ESI-MS/MS analysis of released glycans

Purified glycans were resuspended to a total volume of 9 µL in 0.1% (v/v) TFA in ultrapure water, 8 μL of purified released N-glycans was injected onto a Thermo HyperCarb PGC column (3 μm particle, 1 mm X 30 mm) using an Agilent 1260 HPLC coupled with a Thermo LTQ Velos Pro linear ion trap mass spectrometer. Glycans were chromatographically resolved over 60 min at a flow rate of 15 μL/min at 50 °C with the following gradient: Buffer A: 10 mM ammonium bicarbonate, Buffer B: 70 % (*v/v*) acetonitrile in 10 mM ammonium bicarbonate. The gradient parameters were: 0-3 min—0 % B, 4 min – 14 % B, 40 min – 40 % B, 48 min – 56 % B, 50 - 54 min – 100 % B, 56 – 60 min – 0 % B. The mass spectrometer was operated in negative ion mode and configured to perform one full zoom scan MS experiment (HESI source temperature 55 °C, spray voltage 2.75 kV, sheath gas flow 13, auxiliary gas flow 7, capillary temperature 275 °C, scan range 570-2000 *m/z*, AGC of 3e4, 3 microscans, and maximum IT of 100 msec) with the top 5 precursors (dynamic exclusion window of 15 sec) selected for MS/MS (scan range 150-2000 *m/z*, AGC of 1e4, maximum IT of 100 ms, isolation window 1.4 *m/z*, and normalized collision energy set as 33).

Released glycan analysis was performed manually using the following criteria: *m/z* signals corresponding to biosynthetically possible glycan compositions according to known abundant bovine fetuin N-glycan masses as well as the theoretical products of the enzymes used were selected for area-under-curve (AUC) quantitation using Skyline (26, 27). Each peak area was then expressed as a percentage of the total area of all glycans in the sample, and as a percentage of glycans from its theoretical precursor glycan. Structural characterisation was performed for glycans with MS/MS fragmentation using diagnostic ions (28-30) as well as known PGC elution patterns of specific glycan features (31-33). Two-way ANOVAs comparing time points of each glycan were performed with GraphPad Prism v9.5.1. All raw data is available on Glycopost (34) (Identifier: GPST000738).

## Results

### Initial design and iteration of the IMER housing

Iterative on-column designs were made and tested for the IMER, starting with a v1 prototype of two plastic syringes welded together (Supplementary S1). Following this, an arbitrary internal column volume of 100 µL was chosen for 3D designs and prints of subsequent 3D printed columns (Supplementary S1). Subsequent column versions varied in internal diameter, wall thickness, and connection points, with the final version of the IMER utilising two Luer lock fittings for a stable connection to HPLC and FPLC fittings, as well as thick column walls able to handle measured pressures of up to 140 bar (observed to occur during initial column conditioning, data not shown) without leaking. Ni-NTA resin-bound recombinant glycosyltransferases, immobilised via an attached His-tag, were installed in the IMER, and held in place with a cotton plug (Figure 1) in this column. Each individual IMER was loaded with only one glycosyltransferase.

**Figure 1.**
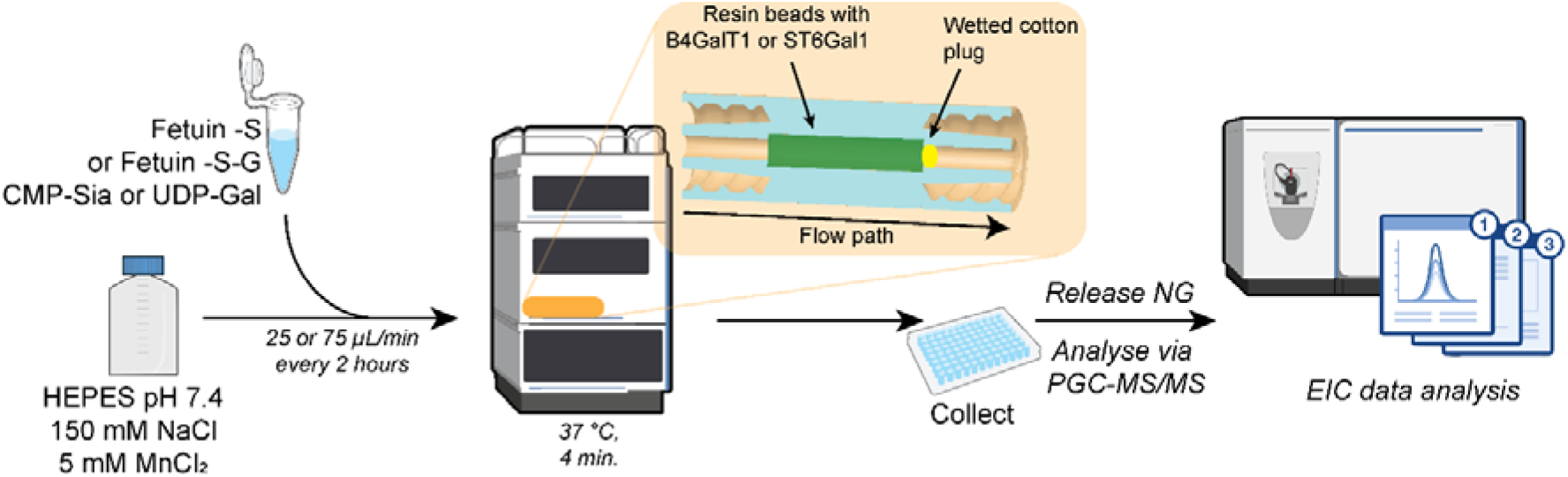
Typical workflow for IMER experimentation. Intact substrate glycoproteins with relevant activated sugar precursors are injected onto a system containing a pre-loaded IMER. Running buffer of the IMER contains essential co-factors such as manganese and is recyclable to minimise buffer usage. After treatment by the IMER, samples are collected and then directly prepared for released N-glycomics analysis via PGC-MS/MS. Injections were performed every 2 hours to test enzyme activity longevity.

Two IMERs were assembled for this work, one containing β-1,4-galactosyltransferase (B4GalT1) and one containing α-2,6-sialyltransferase (ST6Gal1), adding galactose or Neu5Ac residues to their respective substrates. For operation, a packed IMER was installed in the 37 °C column oven of an Agilent 1260 HPLC, with the substrate glycoprotein and activated sugar donor injected over a recyclable running buffer containing all essential co-factors (including manganese) to minimise buffer usage (Figure 1). Substrate was introduced as discrete injections at 0, 2, 4 and 8 hours, with a baseline flow of 5 µL/min maintained between injections, and eluted fractions collected directly into a PVDF membrane plate for released N-glycan analysis by PGC-MS/MS (Figure 1).

Each IMER was operated at two flow rates (25 and 75 µL/min), corresponding to substrate residence times of approximately 240 and 80 seconds respectively, assuming no column retention. As the model substrate, bovine fetuin (Uniprot P12763) carries predominantly tri- and bi-antennary N-glycans (∼85 % and ∼15 % respectively), and the products of each IMER were assessed at both the compositional and isomeric level. Throughout, results are reported as relative family abundance (RFA), the abundance of a glycan as a proportion of its biosynthetic family (a precursor glycan and the products that can be made from it, Supplementary Figure S4), rather than as a proportion of the total N-glycome, as this more directly reflects the precursor-to-product conversion driven by each enzyme.

### Adding galactose residues to precursor glycans with a β-1,4 galactosyltransferase Immobilised enzyme reactor (B4GalT1-IMER)

Terminal galactosylation is a prerequisite for sialylation and other branch elaborations, and the degree of Fc galactosylation modulates effector functions such as complement activation and antibody-dependent cytotoxicity (35). To test the effectiveness of B4GalT1 in the on-column format, bovine fetuin was treated with both sialidase and galactosidase (Fet -S-G) before attempts were made to re-galactosylate the glycans presented on Fet -S-G using immobilised B4GalT1 (B4GalT1-IMER). Some incomplete degalactosylation was observed, indicating a partially heterogeneous starting glycan mixture (Fig. 2B).

**Figure 2.**
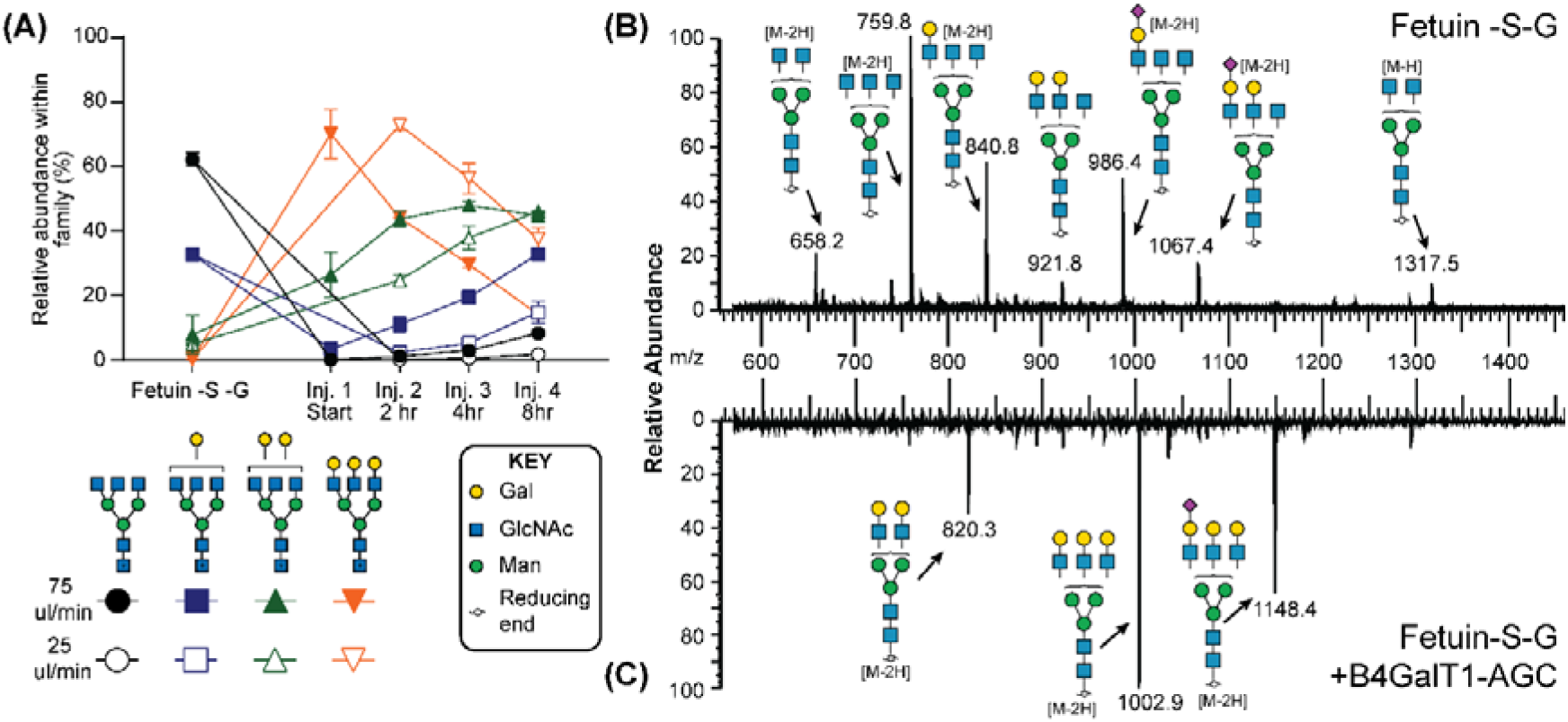
Galactosylation of de-sialylated and de-galactosylated fetuin (Fetuin - S -G) by the B4GalT1-IMER. (A) Relative abundance within the tri-antennary H3N5 glycan family (RFA), measured for the Fetuin -S -G starting control and across four discrete injections (Start, 2, 4 and 8 hours) at two flow rates, with filled symbols denoting 75 µL/min and open symbols 25 µL/min. Inj 1 for 25 μl/min was not included due to loss during poor acquisition quality. (B) Summed released N-glycan spectrum of Fetuin -S -G before treatment. (C) Summed released N-glycan spectrum after treatment with the B4GalT1-IMER.

The B4GalT1-IMER was kept under continuous reaction conditions, and discrete substrate injections were performed at several time points over 8 hours to assess the performance (i.e. longevity of useful activity) of the IMER, starting with an initial injection after 5 minutes of equilibration.

To compare the effectiveness of the B4GalT1-IMER based on residence time, we assessed the conversion of the major tri-antennary substrate to its fully galactosylated form (Figure 2A). The slower flow rate of 25 µL/min resulted in a more complete glycosylation event than at 75 µL/min. The largest difference was observed in H6N5 glycans at the second hour, with RFA of this glycan measured to be 72.9 ± 1.6 %, compared to 44.2 ± 0.1 % found at 75 µL/min (an average difference of 28.7 %, Supplementary Table S1). Partially galactosylated glycans also increased in abundance, with an average difference of 27.6 % RFA between flow rates for H5N5.

After passing through the B4GalT1-IMER, Fetuin-S-G was substantially modified with additional galactose residues on both the tri-and bi-antennary N-glycans, with galactosylated products visible in the released N-glycan spectra before and after treatment (Figure 2B and 2C). Galactosylation was most complete at the first injection, where the tri-antennary precursor H3N5 fell below the limit of detection and the fully galactosylated H6N5 rose from 0.2 ± 0.04 % to 72.9 ± 1.6 % RFA at the 25 µl/min flow rate, shifting the tri-antennary population from a majority H3N5 to a majority H6N5 structure (Figure 2A).

Bi-antennary glycans showed similar results, with the precursor H3N4 glycan dropping in abundance from 65.0 ± 7.4 % to 1.6 ± 2.9 % RFA in the first injection, and complete galactosylation (as shown by the appearance of H5N4) increasing from 3.4 ± 1.1 % to 68.5 ± 1.6 % RFA at the first time point of the 25ul/min flow rate (Supplementary Figure S5, Supplementary Table S1). The effectiveness of galactosylation on injected precursor was found to decrease as the column was kept at reaction conditions over time, with complete galactosylation dropping from 68.5 ±

1.6 % and 72.9 ± 1.6 % RFA in bi- and tri-antennary N-glycans respectively in the first injection to 36.2 ± 5.3 % and 37.5 ± 3.4 % RFA at the fourth sequential injection, 8 hours after the initial injection for the same flow rate. At this time point, partial galactosylation was still observed, via H4N5 and H5N5 glycans.

At the isomeric level, partially galactosylated N-glycans showed a clear preference for specific structural isomers, inferring a preference by B4GalT1 for the addition of galactose to the α,1-6Man arm of the glycan core (Supplementary Figure S6), as determined by MS retention time rules and MS/MS analysis, and matching previously reported preferences of B4GalT1 galactose addition (36).

### Adding Neu5Ac to precursor glycans using an α-2,6 sialyltransferase immobilised enzyme reactor (ST6Gal1-IMER)

Sialic acids are a critical capping sugar, and as such have a great effect on glycoprotein attributes. For antibodies, α2,6-sialylation caps N-glycan antennae and governs serum half-life, immunogenicity and receptor recognition of glycoprotein therapeutics (37). To test the effectiveness of a sialyltransferase IMER (ST6Gal1-IMER), bovine fetuin was treated with α2-3,6,8 Neuraminidase to create a desialylated fetuin (Fet -S) before injection onto the ST6Gal1-IMER. Precursor and product glycans from Fet -S were identified for this IMER (Supplementary Figure S2), and investigated at both a compositional and isomeric level (Figure 3).

**Figure 3.**
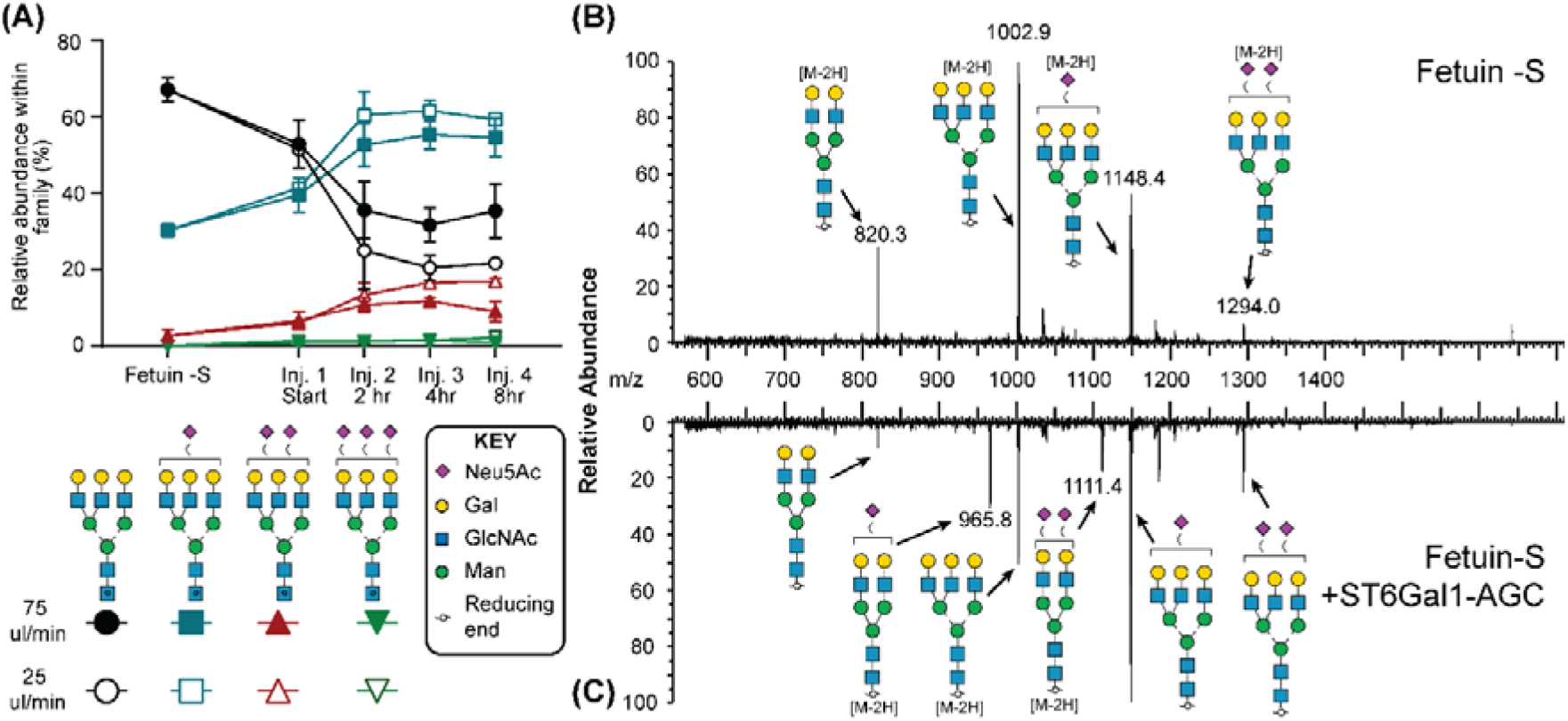
Sialylation of de-sialylated fetuin (Fetuin -S) by the ST6Gal1-IMER. (A) Relative abundance within the glycan family over four injections across 8 hours at two flow rates (25 and 75 µL/min), showing depletion of asialylated precursors and accumulation of sialylated products. (B) Summed released N-glycan spectrum of Fetuin -S before, and after (C) treatment with the ST6Gal1-IMER, showing sialylated products (arrows). Numbers in brackets denote charge state.

Like the B4GalT1-IMER, the ST6Gal1-IMER was maintained at reaction conditions for eight hours, and the ability of the column to effectively sialylate precursor N-glycans conjugated to fetuin was measured through separate injections at multiple time points, with the first injection occurring 5 minutes after column installation to allow for appropriate equilibration. The slower flow rate of 25 µL/min showed a more complete level of sialylation, corresponding to an increase in residence time of the glycoprotein substrate with the enzymes bound to the IMER (Figure 3A). An average difference of 11.9% RFA was observed between flow rates for the initial precursor, H6N5, with 75 µL/min having higher levels of this measured precursor at all time points, and partial sialylation more pronounced at a 25 µL/min flow rate at all time points from the second injection onwards. For this IMER performance assessment, data from both flow rates was combined to increase statistical significance, which results in the sample being in contact with the ST6Gal1-IMER for an average time of 160 seconds.

After flowing through the ST6Gal1-IMER, Fet-S N-glycans were substantially modified by sialylation, producing a markedly different N-glycan population from the injected substrate (Figure 3A), a change also evident in the released N-glycan spectra before and after treatment (Figure 3B and 3C). Unlike the B4GalT1-IMER, which was most active at the first injection, sialylation at the first injection was incomplete, consistent with the column requiring a longer equilibration than the 5 minutes allowed before injection. From the second injection onward, sialylation was consistent across the remaining timepoints (Figure 3A), indicating stable enzyme activity over the 8-hour period. At no point was a significant increase of the triply sialylated product, H6N5S3 observed. At these post-equilibration timepoints, the abundant precursor H6N5 was depleted relative to the pre-injection control, from 67.2 ± 3.1 % to 26.1 ± 7.2 % RFA, with a correlative rise in the partially sialylated tri-antennary products H6N5S1 (30.2 ± 1.7 % to 60.4 ± 0.06 % RFA) and H6N5S2 (2.6 ± 1.7 % to 13.4 ± 2.9 % RFA) (Figure 3A).

A clear increase in sialylation was also observed for other identified product glycans acting as substrates for the ST6Gal1-IMER. The most profound change was observed in biantennary N-glycans, with the third injection resulting in the product N-glycan H5N4 decreasing in RFA from 96.7 ± 1.4 % to 10.6 ± 0.4 % after the column had spent four hours at continuous reaction conditions at 25 µL/min (Supplementary Figure S7). Complete sialylation was observed, with all repeat injections showing a RFA of between 28.9 ± 0.4 % and 40.9 ± 2.2 % for the production of the totally sialylated product glycan, H5N4S2.

At the isomeric level, the approximate ratio of H6N5S2 isomers moved from 4.6 : 3.1 : 12.4 : 10.0 (isomers a, b, c, and d respectively) to an approximate average of 1.4 : 12.7 : 25.1 : 10.0 in ST6Gal1-IMER treated fetuin (all injections combined), showing an increased abundance of isomers with sialic acid on the α-1,3Man arm (isomers b, c, and d, Supplementary Figure S8) rather than on the α-1,6Man arm of the N-glycan core (isomer a), consistent with the reported preference of ST6Gal1.

## Discussion

Immobilised enzyme reactors (IMERs) are well established for biopolymer degradation, proteomics, biomarker discovery, inhibitor screening, and detection (19, 38), but their use for the addition of sugars to substrates remains limited. Existing examples include a magnetic nanoparticle–based ‘Golgi-on-a-chip’ (39), and a monolithic oxazoline IMER in which engineered EndoS variants with glycosidase and transglycosidase activity transfer complex oxazoline glycans to an on-column de-glycosylated IgG antibody (40). More recently, solid-phase and flow-based glycan remodelling platforms have been developed. The SUGAR-TARGET platform (18) uses glycosyltransferases immobilised on beads via a biotin-streptavidin linkage to achieve efficient galactosylation of IgGs (80.2–96.3 % terminal galactosylation), but operates as a series of discrete batch reactions in which the glycoprotein target is manually separated from the IMER between steps rather than as a continuous flow-through system. The microgel-based MiRAGE reactor (21) similarly couples immobilised glycosyltransferases, here in a continuous membrane reactor with *in situ* removal of low-molecular-weight by-products, but is directed at the de novo assembly of free oligosaccharides (such as human milk oligosaccharides and blood-group epitopes) rather than the modification of glycans already present on a purified glycoprotein. In contrast, the IMER employed here is a continuous flow-through column that acts directly on the existing N-glycans on an intact, purified glycoprotein, in a hands-off format that can be applied to successive substrates without an interstitial purification or enzyme-removal step. With the right choice of enzymes it is conceivable that this IMER is also suitable for other glycan types, such as O-glycans.

IMERs create a local environment where immobilised enzyme concentration is much higher than would typically occur in a one pot reaction, leading to rate-limited rather than enzyme-limited glycosylation reactions. Both IMERs tested achieved significant galactosylation and sialylation of fetuin within contact times of up to four minutes, resulting in a distinctly different glycoprofile on the native protein. These same enzymatic reactions are typically performed overnight in a one-pot vessel (41), so this represents a substantial time saving even where reactions do not currently reach 100 % completeness. The trade-off between contact time and conversion observed here is consistent with other continuous-flow systems, where short substrate residence times have likewise been shown to limit conversion relative to extended one-pot reactions (21). Completeness could be improved by utilising faster-acting enzymes (42), engineering existing enzymes to react faster (43, 44), or modifying the physical parameters of the column itself. For industrial control of glycosylation as a CQA, total reaction completeness may be required to minimise heterogeneity, though an activity-biased glycomic profile may be sufficient for improved product synthesis.

By separating glycosyltransferases into individual chambers, as used here, there is potential for ‘dial-up’ sequential glycan addition, where each column presents the precursor required by the next. Connecting IMER chambers in sequence would create a plug-and-play system in which a defined protein glycan can be built from a chosen array of glycosyltransferases, an arrangement that parallels the sequential modular reactors recently realised under continuous flow in the MiRAGE platform (21) and the discrete bead-based steps of SUGAR-TARGET (18).

Isomeric arm-addition preferences were analysed for both enzymes. B4GalT1 showed a marked preference for the α,1-6Man arm of the glycan core (Supplementary Figure S6), consistent with the 1,6-arm-first galactosylation reported for immobilised GalT elsewhere (36), and a preference that could be exploited to direct arm-specific glycoengineering. Despite this bias, complete galactosylation was still achieved, confirming B4GalT1 as a robust enzyme with broad substrate specificity across the available N-acetylglucosamine-terminating arms. Similarly, the reported preference of ST6Gal1 for sialic acid addition on the α,1-3Man branch of a biantennary N-glycan (37, 45) was observed in the isomers of the tri-antennary N-glycan H6N5S2 (Supplementary Figure S8), although the reaction began from a skewed population of precursor isomers, which may have biased the observed isomeric ratios. Controlled sialylation and its linkage are biologically important, as sialylation patterns affect how a glycoconjugate interacts with its surrounding environment, and have been well documented for glycoproteins such as IgG (46) and EPO (47). The ability of the IMER to impose arm-specific sialylation bias, alongside the complete galactosylation seen with B4GalT1, demonstrates that immobilised glycosyltransferase chambers can both drive reactions to high conversion and exploit intrinsic enzyme regiochemistry for targeted glycoengineering.

Here we demonstrate a proof-of-concept immobilised enzyme reactor for directed post-production glycan modification of purified glycoproteins under physiological conditions. Both the B4GalT1-IMER and the ST6Gal1-IMER effectively added their respective monosaccharides to pre-treated bovine fetuin N-glycans within an average substrate-enzyme contact time of ∼160 seconds, achieving near-complete galactosylation and substantial α2,6-sialylation, and isomeric-level analysis showed that intrinsic enzyme arm-preferences were retained in the immobilised format. The IMER therefore does not compromise enzyme regio-selectivity while enabling rapid, on-column glycan extension.

By performing *in vitro* glycosylation in IMERs, the complexity of glycosylation within the host cell is converted into controllable parameters that can be optimised and engineered. Each physical parameter of the IMER (temperature, flow rate, substrate amount, column size, and the method of enzyme immobilisation) can be tuned independently and even the limited range of flow rates tested here produced a marked effect on reaction completeness. This suggests that further optimisation of these parameters alone could bring the IMER close to quantitative conversion if this is a requirement. Equally, the mammalian-expressed human glycosyltransferases used here could be replaced with more readily produced bacterial or yeast equivalents (e.g. LgtB from *Neisseria meningitidis* (48) for β1,4-galactosyltransferase activity, or Pd26ST from *Photobacterium damselae* (49) for α2,6-sialyltransferase activity), reducing upstream cost and complexity. The broader catalogue of bacterial glycosyltransferases covering α2,3-sialylation (50), core fucosylation (51) and other transferase activities would also expand the range of defined glycan structures accessible through this platform.

Together, these findings establish the IMER as a viable approach to post-production glycan finishing. Glycoproteins with high native glycan heterogeneity could first be specifically deglycosylated, either enzymatically or through *in vivo* glycoengineering (52), then re-glycosylated in a controlled manner by a sequence of IMERs to yield a defined target glycoform. The same approach could instead start from a host that already installs simple or partially complete glycans, such as insect, plant or glycoengineered yeast (7); the truncated, GlcNAc- or mannose-terminating structures these systems produce are well matched to the IMER concept demonstrated here, which extend compatible precursors without prior trimming steps. Such a platform would decouple glycosylation from the chosen protein expression host, allowing production systems that are otherwise attractive in terms of titre, growth rate, or cost, but currently fall short on optimal glycan structure, to be used for the generation of recombinant glycoprotein therapeutics with controlled glycosylation as a critical quality attribute.

## Author Contributions

ND and EM contributed to experimentation, idea generation, data analysis, editing, and proofing. JC, CL contributed to experimentation and data analysis. NP contributed to supervision, editing, proofing, and funding.

## Acknowledgements

The authors wish to acknowledge the contributions of Jack Rowlatt, Andrea Maggioni and Daniel Kolarich for their support. The authors wish to acknowledge the plasmids from the Complex Carbohydrate Research Center (glycoenzymes.ccrc.uga.edu) and the grant support for the Repository (National institutes of Health grants P41GM103390 and P01GM107012).

## Supplementary material

**Table S1.**
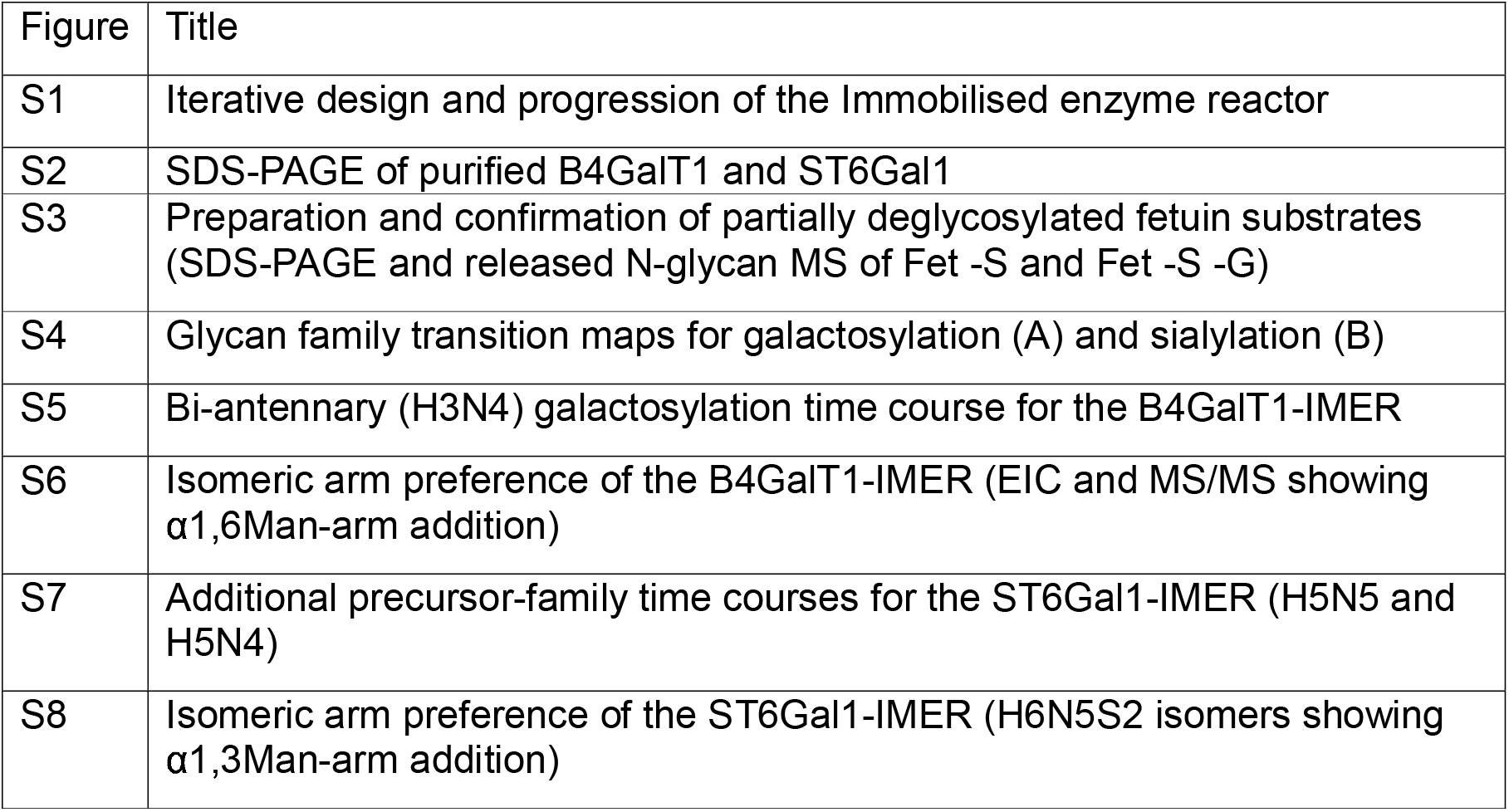
Raw data of measured relative abundance of glycans.

