## Supplementary_File_1 for "Immobilised enzyme reactors for post-production glycan modification of purified glycoproteins"

***Correspondence:**
Dr Edward S.X. Moh

**Supplementary material**

| Figure | Title |
| --- | --- |
| S1 | Iterative design and progression of the Immobilised enzyme reactor |
| S2 | SDS-PAGE of purified B4GalT1 and ST6Gal1 |
| S3 | Preparation and confirmation of partially deglycosylated fetuin substrates (SDS-PAGE and released N-glycan MS of Fet -S and Fet -S -G) |
| S4 | Glycan family transition maps for galactosylation (A) and sialylation (B) |
| S5 | Bi-antennary (H3N4) galactosylation time course for the B4GalT1-IMER |
| S6 | Isomeric arm preference of the B4GalT1-IMER (EIC and MS/MS showing α1,6Man-arm addition) |
| S7 | Additional precursor-family time courses for the ST6Gal1-IMER (H5N5 and H5N4) |
| S8 | Isomeric arm preference of the ST6Gal1-IMER (H6N5S2 isomers showing α1,3Man-arm addition) |

**Figure S1 -** Iterative design and progression of the Artificial Golgi Column

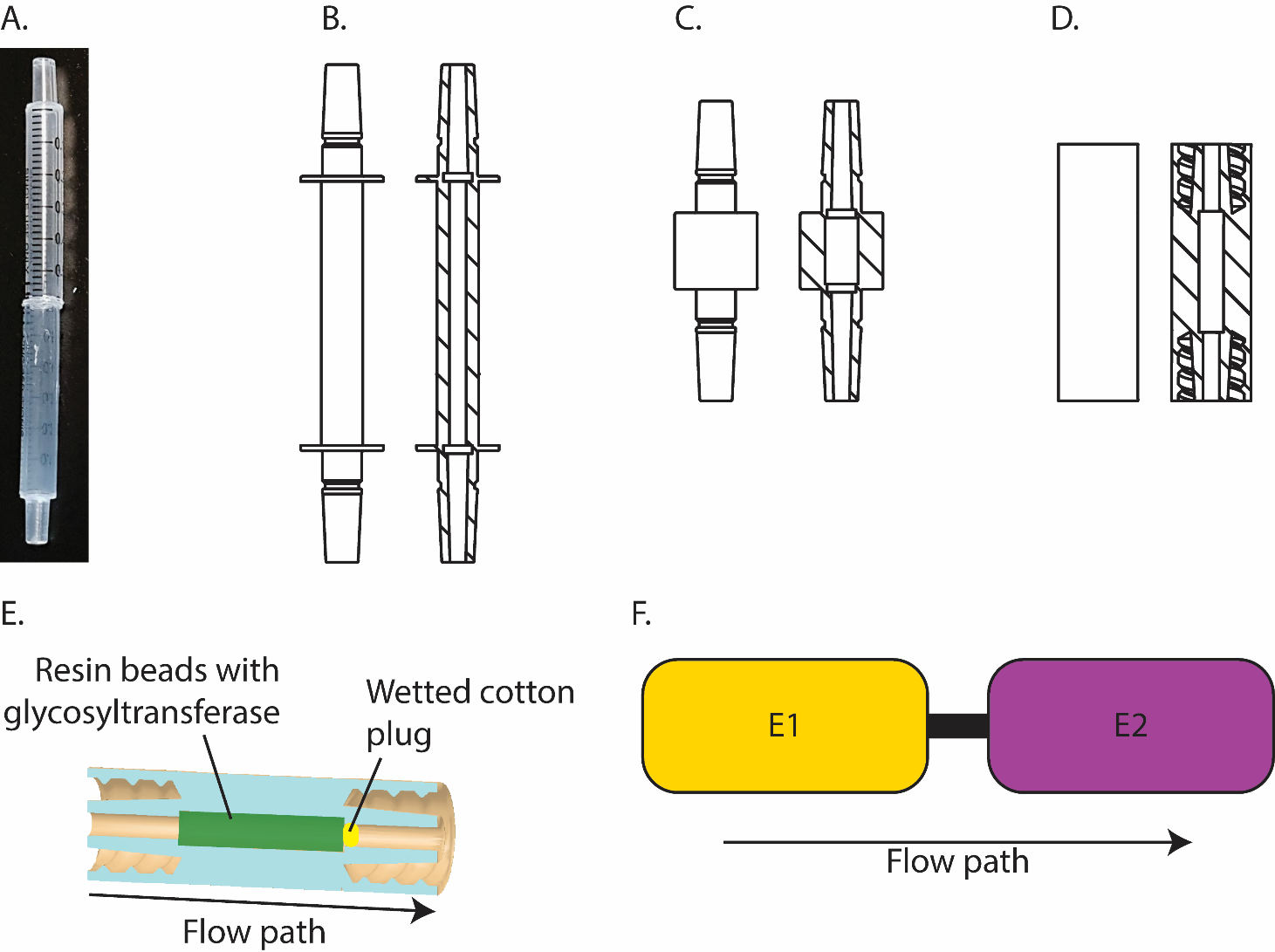

Iterative designs of the Artificial Golgi Column (IMER) housing. A. Initially IMER columns were made using two 100 µl disposable syringes welded together, then columns were 3D printed with specific dimensions. B. The second iteration of columns had a 2 mm internal diameter (ID) and a luer slip fitting. C. The third iteration of IMERs moved to a thicker column wall and wider (4 mm) ID. D. Final IMER column housings employed a more secure luer lock fitting rather than luer slip while retaining the 4 mm ID. Models B, C and D are to scale with each other.

**Figure S2 -** SDS-PAGE of purified β4GalT1 and ST6Gal1

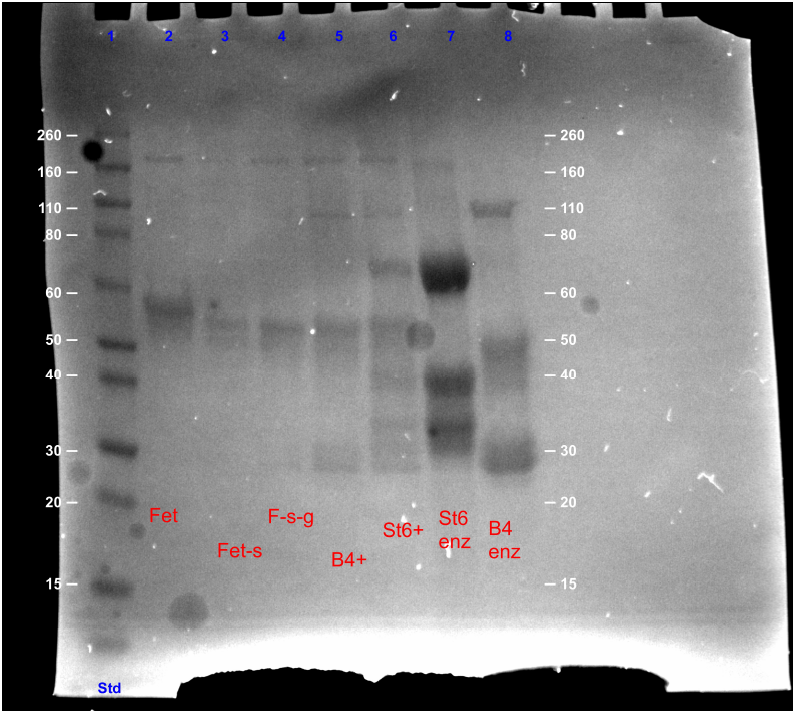

SDS-page gel showing fetuin pre (L2) and post treatment with sialidase (L3), and sialidase and galactosidase (L4). Fet-S-G after treatment with B4GalT1 (L5) and Fet-S after treatment with ST6Gal1 (L6) in solution is also shown. Finally, Lanes 7 and 8 show the purified ST6 and B4GalT1 enzymes respectively.

**Figure S3 -** Preparation and confirmation of partially deglycosylated fetuin substrates SDS-PAGE

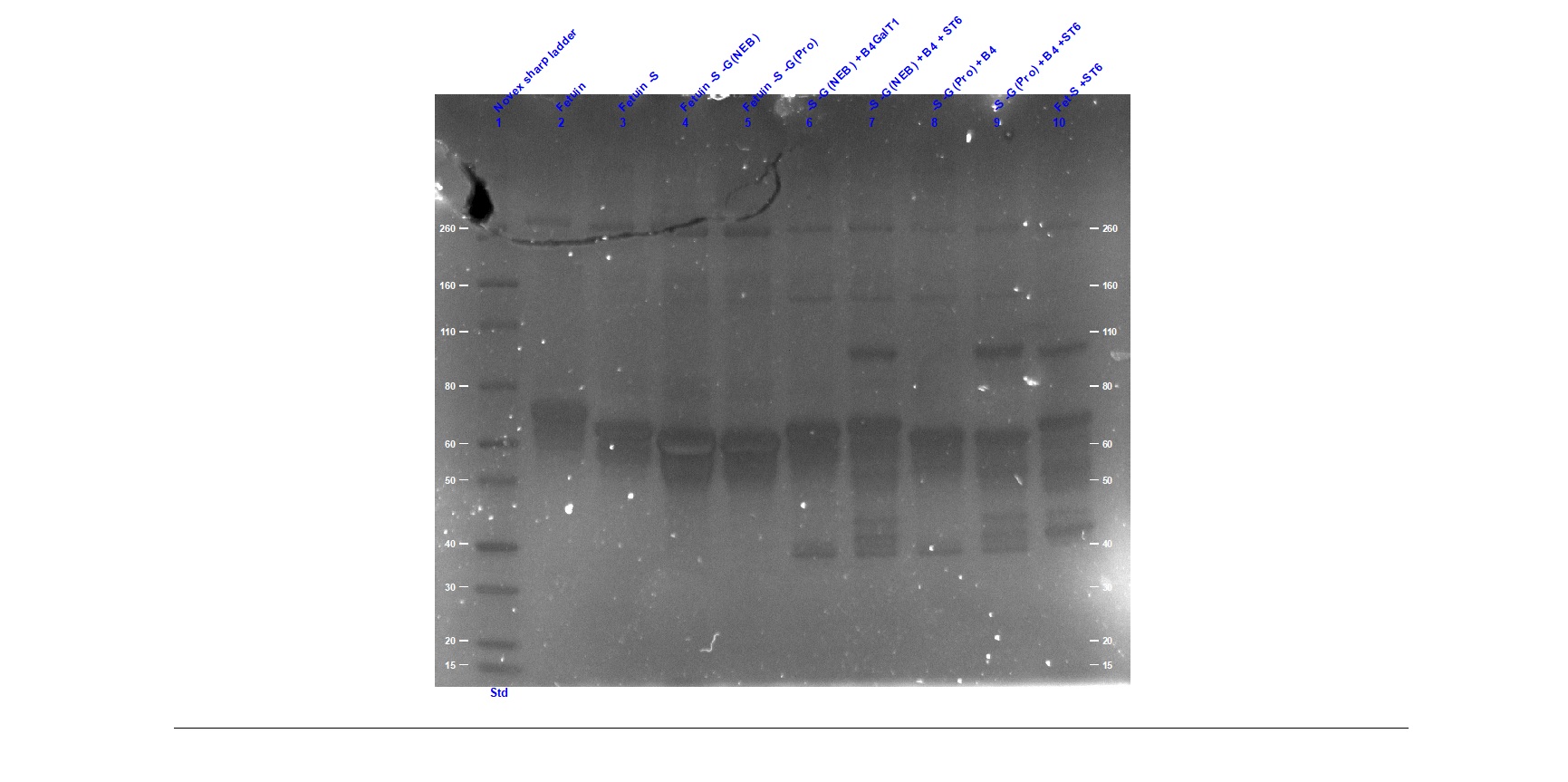

Fetuin after treatment with different sialidases and galactosidases, and then after in solution treatment with both B4GalT1 and ST6Gal1 in solution is shown.

**Figure S4 -** Glycan family transition maps for galactosylation and sialylation

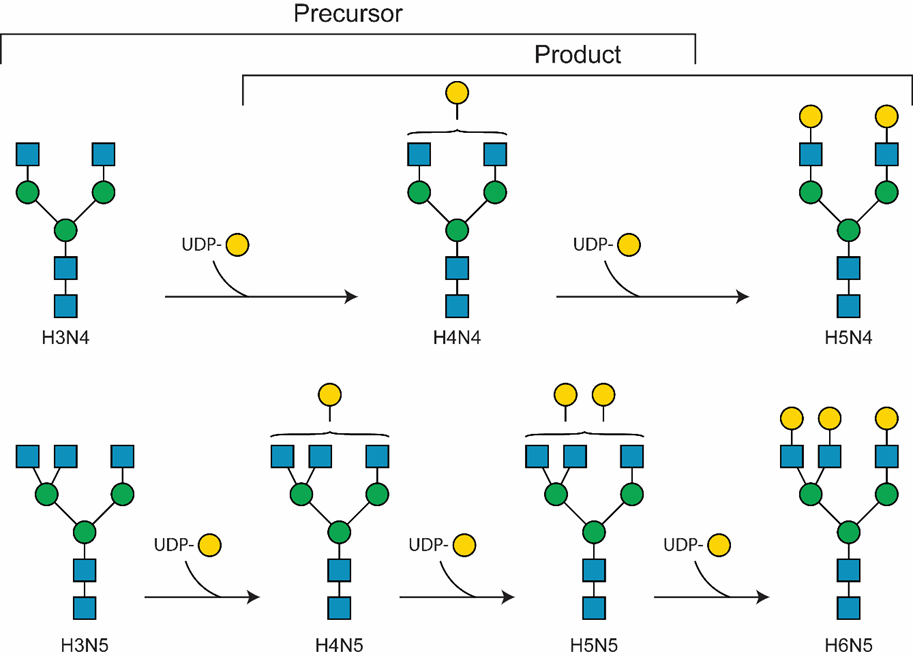

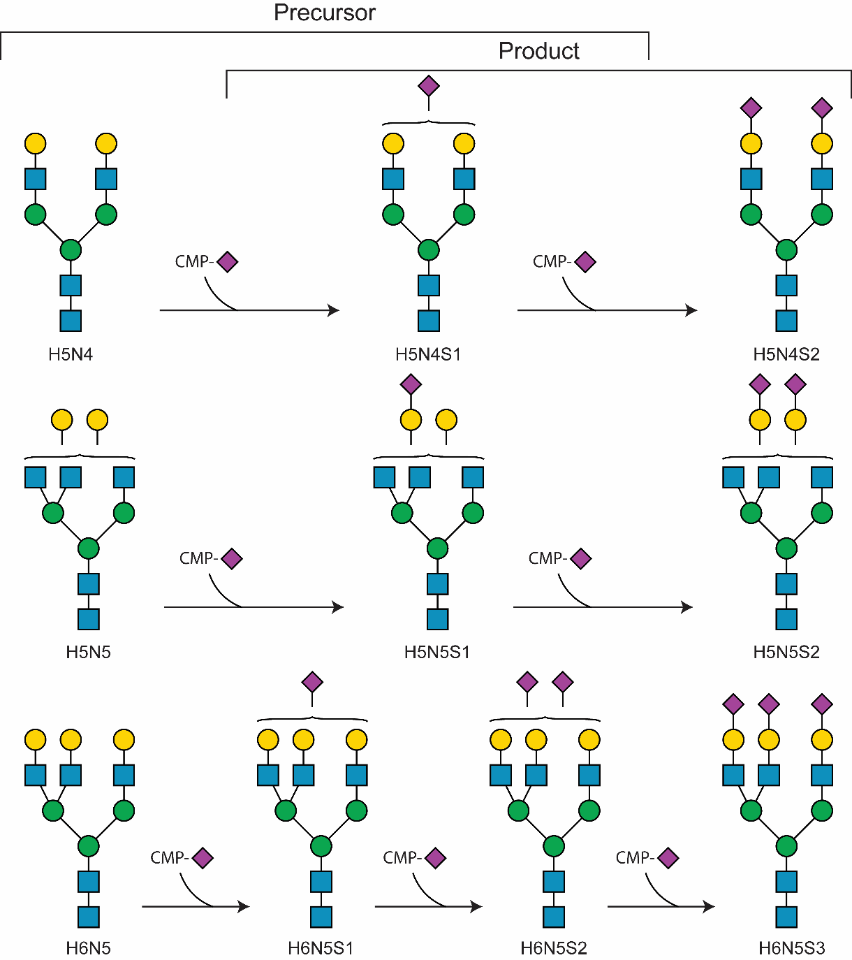

Precursor and product glycan families assessed from foetal bovine serum fetuin deglycosylated with sialidase and galactosidase. Note, that as galactose residues are added, product glycans can also act as precursors for the next galactose addition—tri-antennary N-glycans comprised over 85 % of all glycan abundances produced Product and precursor glycan families assessed from foetal bovine serum fetuin treated with sialidase. Tri-antennary N-glycans comprised over 85 % of all glycan family abundances, and all precursor glycans were present as substrates for subsequent ST6Gal1-IMER experiments.

**Figure S5 -** Bi-antennary (H3N4) galactosylation time course for the β4GalT1-IMER

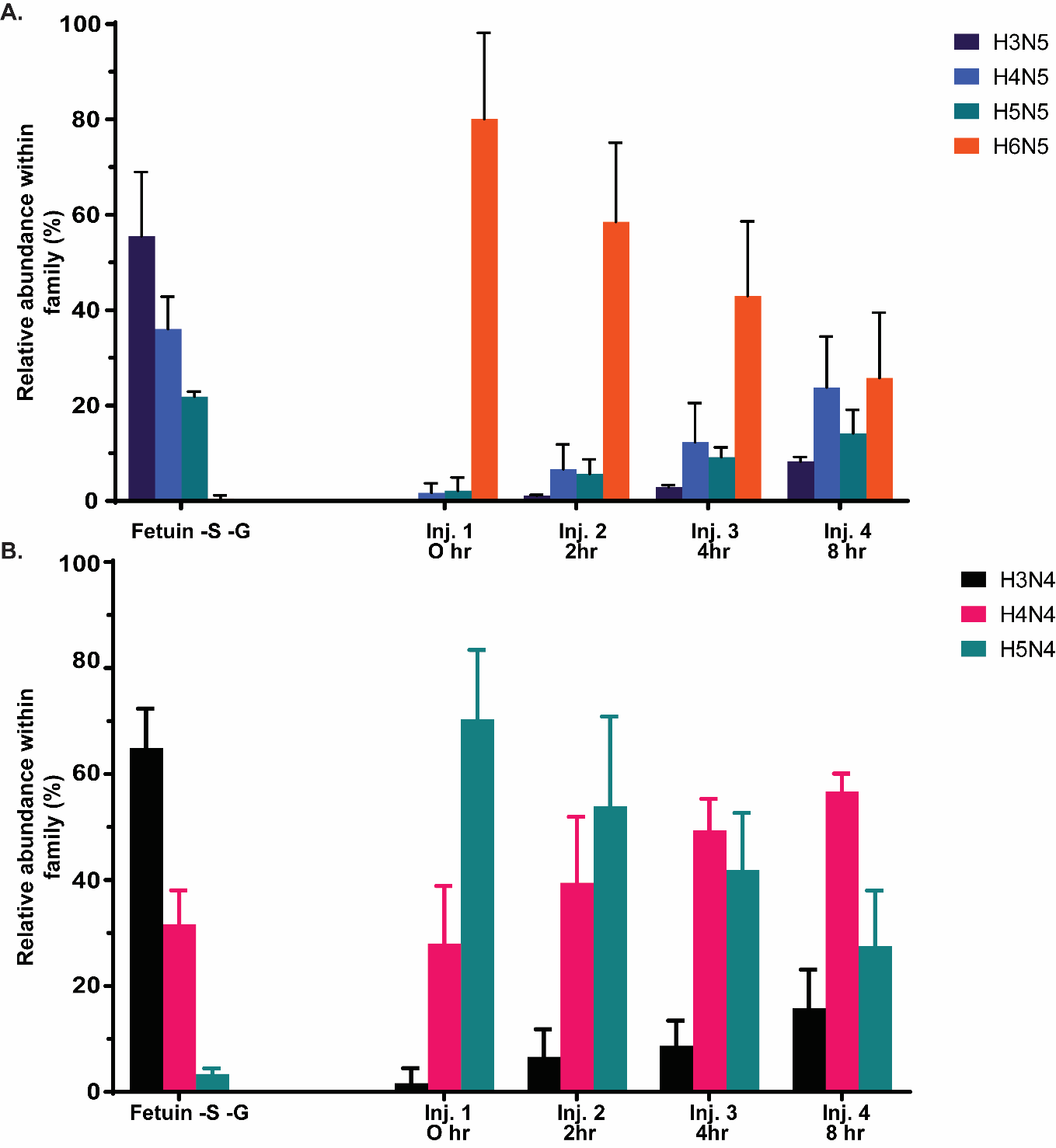

Relative glycan abundance within the glycan family using H3N4 as a precursor glycan. Depletion of non-galactosylated precursor H3N4 is observed at all time points, with subsequent intermediary (H4N4) and complete galactosylation observed (H5N4).

**Figure S6 -** Isomeric arm preference of the β4GalT1-IMER (EIC and MS/MS showing α1,6Man-arm addition)

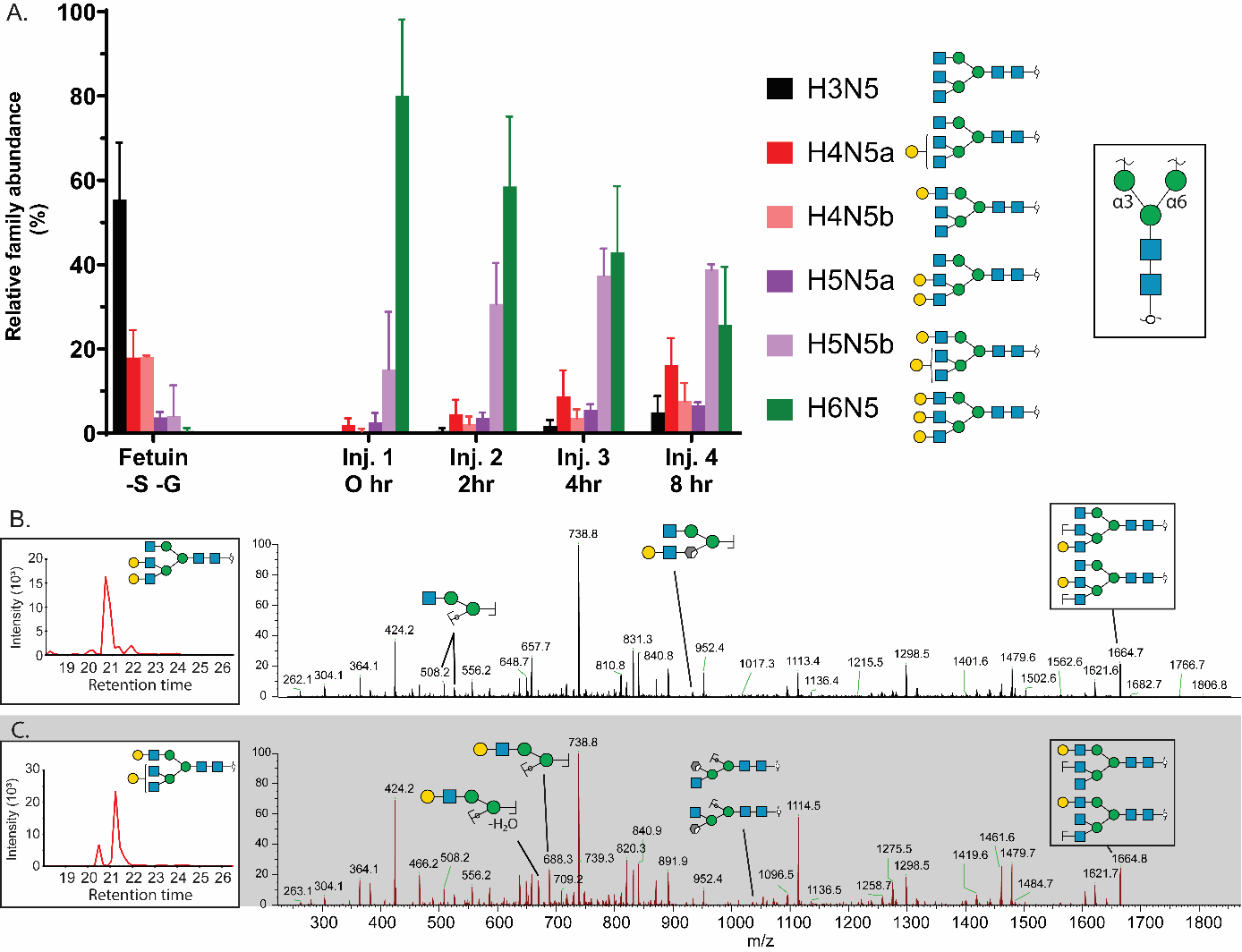

A. Tri antennary isomer differences within one glycan family. A clear preference towards the α-1,6Man arm (top arm) galactosylated glycans can be seen. B. H5N5 glycan EIC before treatment with the β4GalT1-IMER, with accompanying negative mode MS/MS spectra of the most abundant isomer. D and D-18 ions can be observed, showing no galactosylation on the 1,6-arm of this isomer. C. H5N5 glycan EIC after treatment with the β4GalT1-IMER, with accompanying negative mode MS/MS spectra of the most abundant isomer. Here, the D and D-18 ions for 1,6-arm galactosylation can be observed, which combined with the EIC show a clear arm preference for galactose addition.

**Figure S7 -** Additional precursor-family time courses for the ST6Gal1-IMER (H5N5 and H5N4)

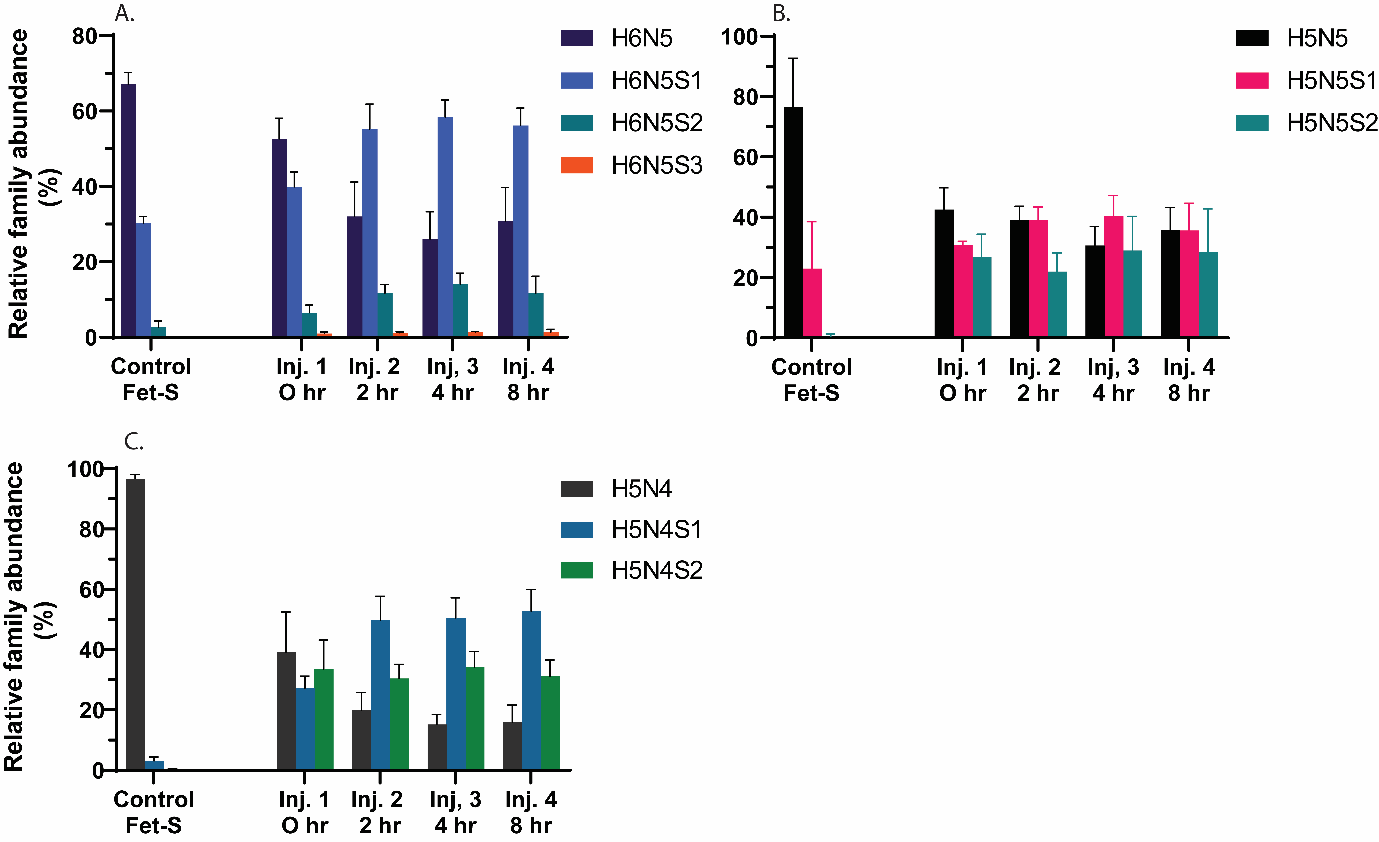

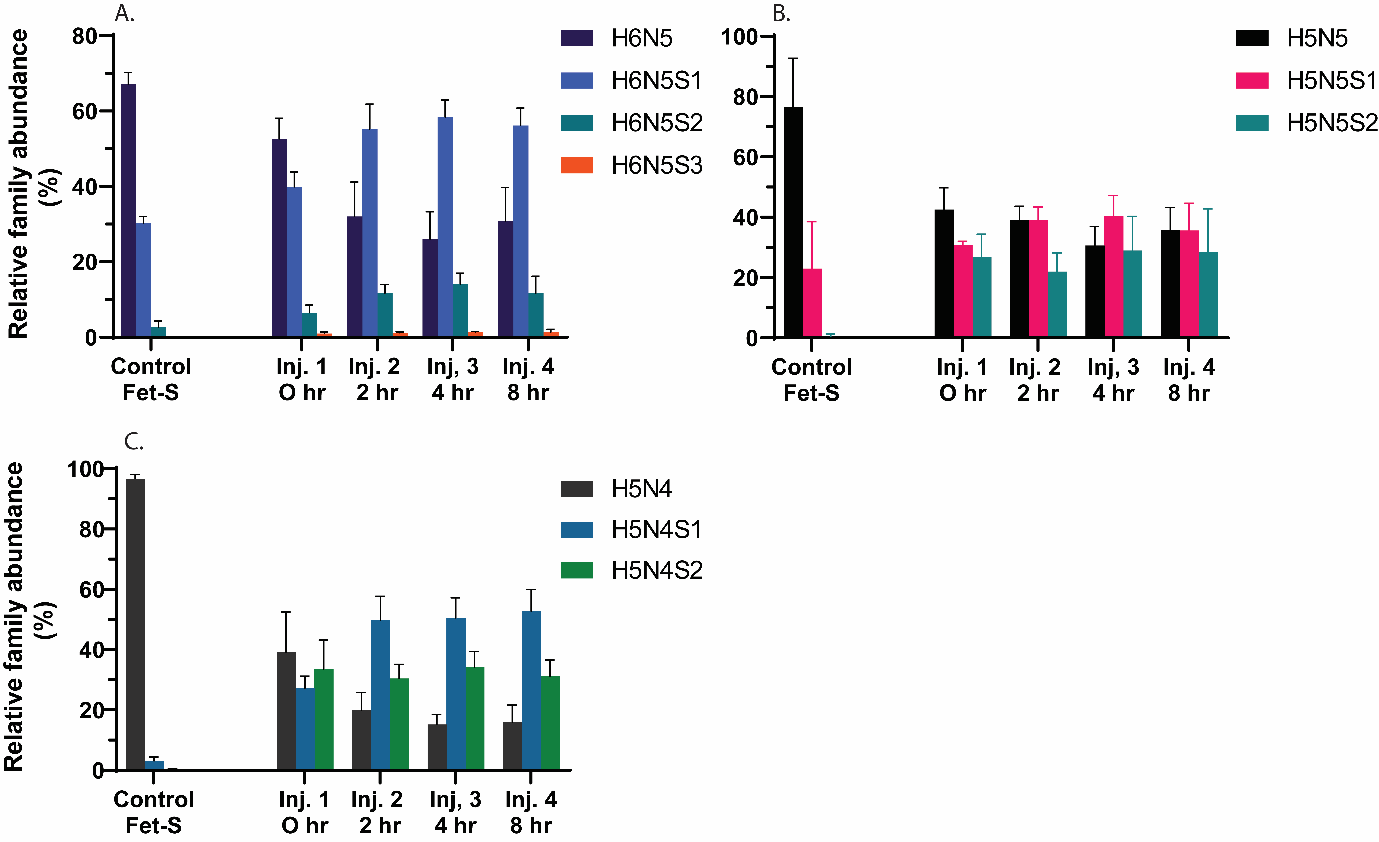

Relative abundance within glycan families of the two observed depleted precursor glycans, H5N5, and H5N4. A significant change in family abundance composition can be observed for all precursor glycans from starting injection, with the de-sialylated glycans significantly reducing, in relative abundance.

**Figure S8 -** Isomeric arm preference of the ST6Gal1-IMER (H6N5S2 isomers showing α1,3Man-arm addition)

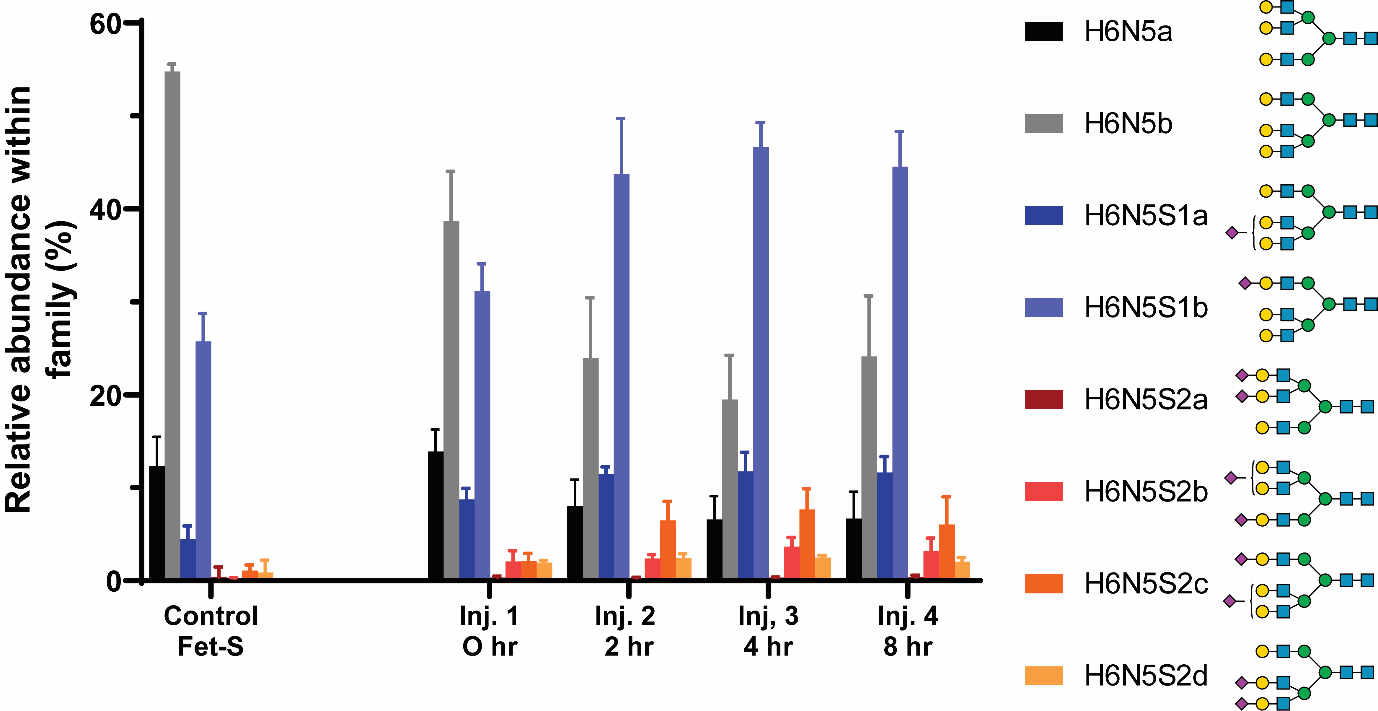

Isomeric differences of H6N5 glycan family. Differences in glycan ratios can be observed between the control desialylated fetuin and each subsequent injection, with a preference for sialic acids to be added to the α-1,3Man (shown at bottom) arm of the N-glycan core.
